# Diverse functions associate with trans-species polymorphisms in humans

**DOI:** 10.1101/2021.01.21.427090

**Authors:** Keila Velázquez-Arcelay, Mary Lauren Benton, John A. Capra

## Abstract

Long-term balancing selection (LTBS) can maintain allelic variation at a locus over millions of years and through speciation events. Variants shared between species, hereafter “trans-species polymorphisms” (TSPs), often result from LTBS due to host-pathogen interactions. For instance, the major histocompatibility complex (MHC) locus contains TSPs present across primates. Several hundred TSPs have been identified in humans and chimpanzees; however, because many are in non-coding regions of the genome, the functions and adaptive roles for most TSPs remain unknown. We integrated diverse genomic annotations to explore the functions of 125 previously identified non-coding TSPs that are likely under LTBS since the common ancestor of humans and chimpanzees. We analyzed genome-wide functional assays, expression quantitative trait loci (eQTL), genome-wide association studies (GWAS), and phenome-wide association studies (PheWAS). We identify functional annotations for 119 TSP regions, including 71 with evidence of gene regulatory function from GTEx or genome-wide functional genomics data and 21 with evidence of trait association from GWAS and PheWAS. TSPs in humans associate with many immune system phenotypes, including response to pathogens, but we also find associations with a range of other phenotypes, including body mass, alcohol intake, urate levels, chronotype, and risk-taking behavior. The diversity of traits associated with non-coding human TSPs suggest that functions beyond the immune system are often subject to LTBS. Furthermore, several of these trait associations provide support and candidate genetic loci for previous hypothesis about behavioral diversity in great ape populations, such as the importance of variation in sleep cycles and risk sensitivity.

**Significance statement:** Most genetic variants present in human populations are young (<100,000 years old); however, a few hundred are millions of years old with origins before the divergence of humans and chimpanzees. These trans-species polymorphisms (TSPs) were likely maintained by balancing selection—evolutionary pressure to maintain genetic diversity at a locus. However, the functions driving this selection, especially for non-coding TSPs, are largely unknown. We integrate genome-wide annotation strategies to demonstrate TSP associations with immune system function, behavior (addition, cognition, risky behavior), uric acid metabolism, and many other phenotypes. These results substantially expand our understanding of functions TSPs and suggest a substantial role for balancing selection beyond the immune system.

## Introduction

The interaction between populations and environments is dynamic. Over time, allele frequencies in a population shift due to drift and adaptive responses to specific environmental pressures. Most genetic variants are short-lived compared to the timescale of species. But on rare occasions variants persistently segregate at intermediate frequencies for millions of years, sometimes predating the most recent common ancestor (MRCA) between two sister species (Bitarello et al., 2018; Cheng & DeGiorgio, 2019; DeGiorgio, Lohmueller, & Nielsen, 2014; Leffler et al., 2013; Siewert & Voight, 2017; Teixeira et al., 2015). These trans-species polymorphisms (TSPs) are likely a sign of genomic regions under long-term balancing selection (LTBS). Over time, instances of LTBS leave signatures in the genome that differentiate them from those under other forms of selection (Bitarello et al., 2018; Key, Teixeira, de Filippo, & Andrés, 2014; Leffler et al., 2013; Siewert & Voight, 2017).

Several instances of likely LTBS have been observed in humans and other primates, mostly within the major histocompatibility complex (MHC) or the ABO blood group locus. For example, the MHC, or human leukocyte antigen (HLA) system in humans, is a family of varied proteins expressed on the cell surface with essential functions in adaptive immune response and regulation. Balancing selection on different components of the HLA region dates to the common ancestor between chimpanzees and humans (Azevedo, Serrano, Amorim, & Cooper, 2015). Similarly, the ABO gene has three alleles, and its variants lead to different blood cell antigens, or lack of thereof, on the surface of the cell. Variation in this group could have a benefit in the immune response to pathogens, and balanced polymorphisms at this locus date back to the common ancestor of gorillas, orangutans, and humans (Ségurel et al., 2012). Other immune-related genes show LTBS between humans and other primates, e.g.: *TRIM5*, a RING finger protein 88 (Battivelli et al., 2011; Cagliani et al., 2010; Ganser-Pornillos & Pornillos, 2019), and *ZC3HAV1*, a zinc finger CCCH-type antiviral protein 1 (Cagliani et al., 2012; De Filippo et al., 2016; Mao et al., 2013; Todorova, Bock, & Chang, 2015). These genes have important roles in host/pathogen response through inhibition of virus replication.

The high allelic variation induced and maintained by balancing selection at certain loci can also be beneficial when a population needs to adapt to new environments. Some variants found under balancing selection in Africa have come to be under directional selection in non-African populations (European and Asian), with one allele becoming predominant in the population (De Filippo et al., 2016). This suggests the adaptive potential of the variation maintained under balancing selection; however, most of these variants were not likely under LTBS.

Recent studies have developed tools and methods to identify instances of balancing selection in genome-wide data (Siewert & Voight, 2017, 2020). Leffler et al. (2013) compared polymorphisms across the genome in Yoruba individuals from the 1000 Genomes Project to those found in Western chimpanzees sequenced by the PanMap Project. They identified coding variants in 324 genes and 125 non-coding haplotypes with multiple TSPs suggesting the presence of LTBS.

Despite the importance and prevalence of balancing selection, most regions bearing signatures of LTBS, including these TSPs, have not been functionally characterized. Determining the functional roles of these polymorphisms in human adaptation and health will deepen our understanding of the dynamics of balancing and positive selection and their roles in adaptation to new environments. Here, we focus on the non-coding TSPs identified by Leffler et al. (2013). We identify potential functional drivers of the LTBS in humans by applying several genome-wide functional annotations and association tests to these TSPs. Our results identify diverse functions, including effects unrelated to the immune system, that may underlie LTBS on the human and chimpanzee lineages.

## Results

### Human-chimpanzee TSPs

We consider 125 human genomic regions containing multiple TSPs that are segregating in both humans and chimpanzees. This set is composed of 263 variants with strong evidence of identity-by-descent; i.e., the divergence between the ancestral and derived alleles is deeper than the human-chimp speciation event. These TSPs were identified by Leffler et al. (2013) based on the observation from coalescent theory (Ségurel et al., 2012) that pairs of TSPs within 4 kb in the human genome are extremely unlikely to result from neutral processes, and thus are strong candidates for LTBS (Figure 1). Hereafter, we refer to these as “TSP regions”.

**Figure 1.**
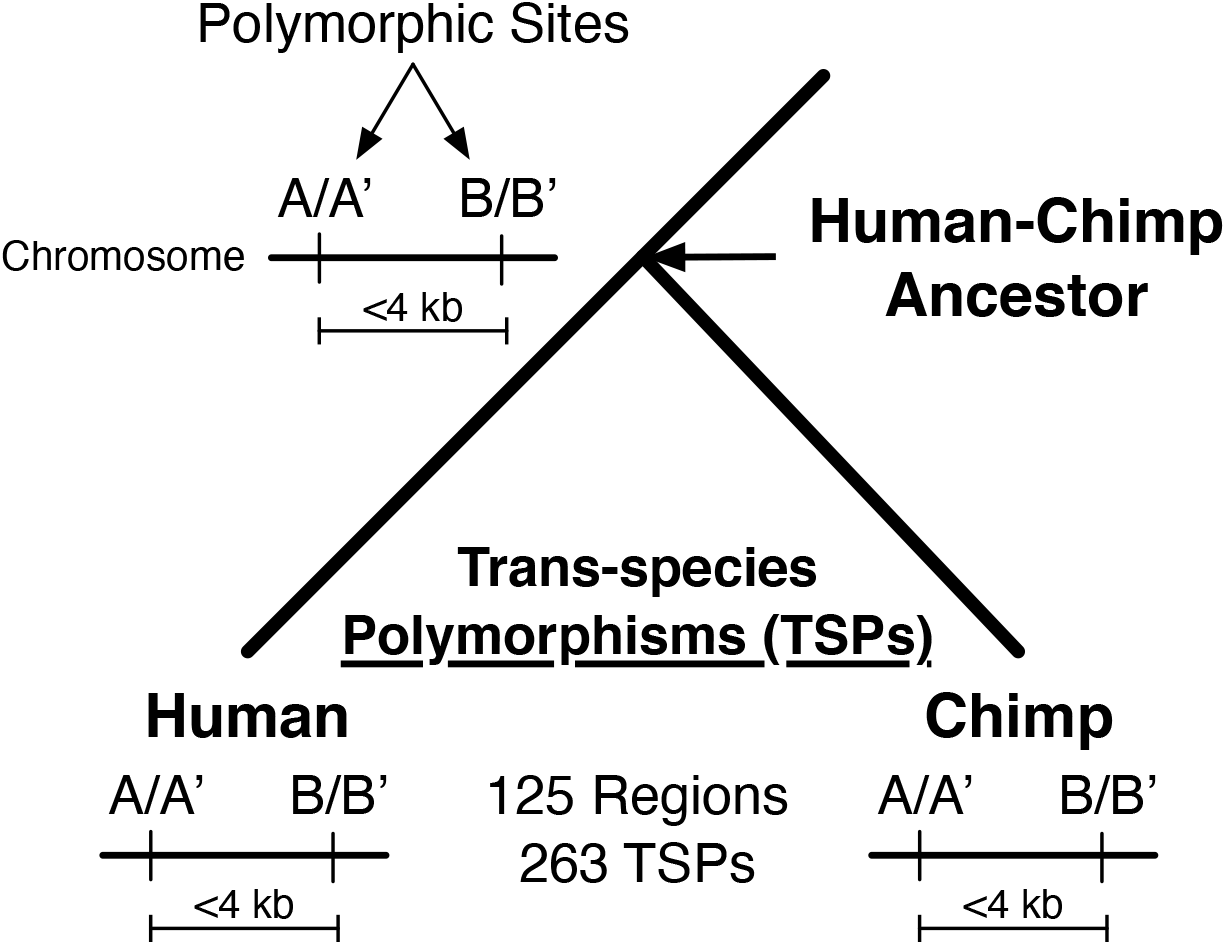
Trans-species polymorphisms (TSPs) likely resulting from long-term balancing selection (LTBS). Schematic showing the criteria used by Leffler et al. (2013) to identify TSPs likely maintained by LTBS. Each line represents a chromosome with polymorphisms segregating in a population. A/A’ are two alleles segregating in both humans and chimpanzees at one site (i.e., a TSP), and B/B’ are alleles segregating in both species at a nearby site. TSPs are very unlikely to appear nearby (within 4 kb) without the action of balancing selection. We consider 125 TSP regions containing 263 TSPs. Within these regions, multiple functional scenarios are possible. For example, one TSP may be under LTBS while the other is neutral, but maintained due to tight linkage. Alternatively, the TSPs may have epistatic functions and both be under selection. In addition to the 263 TSPs, we also considered functional associations with 9,996 variants in high LD (r^2^ > 0.8) with a TSP at least one population from the 1000 Genomes Project (Supplementary Figure 1).

In the following, we analyze two sets of variants for the 125 TSP regions. First, we focus on the 263 TSPs themselves. The average distance between TSPs defining a region is 1652.7 bp (standard deviation = 1303.2). Second, to capture functions tagged by variants in high linkage disequilibrium (LD) with TSPs, we also considered potential tag SNPs in high LD (R^2^≥0.8) with TSPs in African, European, and East Asian populations from the 1000 Genomes Project. This LD-expanded set includes 10,259 variants across the 125 TSP regions (Supplementary Figure 1). By expanding to include LD, we capture additional associations, but may also introduce false positives; thus, we report results on both sets throughout.

### Trans-species polymorphisms overlap diverse functional annotations

We intersected the TSPs with diverse lines of functional evidence from large-scale genomic studies, including genome-wide functional genomics assays, eQTL, GWAS, and PheWAS. We found at least one functional annotation for 95% of the TSP regions (119 out of 125) covering 130 TSPs and 4,807 LD SNPs (Figure 2). Here, we provide an overview of the overlap with these annotations. In future sections, we provide details about each of these annotations. Given that these regions were filtered by Leffler et al. (2013) to exclude coding TSPs, variants in 91% (114 out of 125) of regions overlap annotated gene regulatory regions. This includes 58 TSPs in 40 regions and 1334 LD variants in 112 regions. We also found 86 TSPs across 51 regions with evidence of being expression quantitative trait loci (eQTL) in 29 tissues. Including the variants in LD with TSPs, 57.6% of the regions (72 out of 125) contain eQTLs (p < E-5) in 49 tissues. We found genome-wide significant associations with phenotypes in available genome- or phenome-wide association studies for 19% of the regions (24 out of 125; 13 GWAS and 16 geneAtlas PheWAS). Finally, 11.2% (14 out of 125) of these regions contain SNPs in protein-coding regions; two TSPs in different regions produce non-synonymous protein sequence changes, and 17 other non-synonymous variants in 7 regions are in high LD with the TSPs. (These were not identified as coding in the previous study due to changes in protein annotations.)

**Figure 2.**
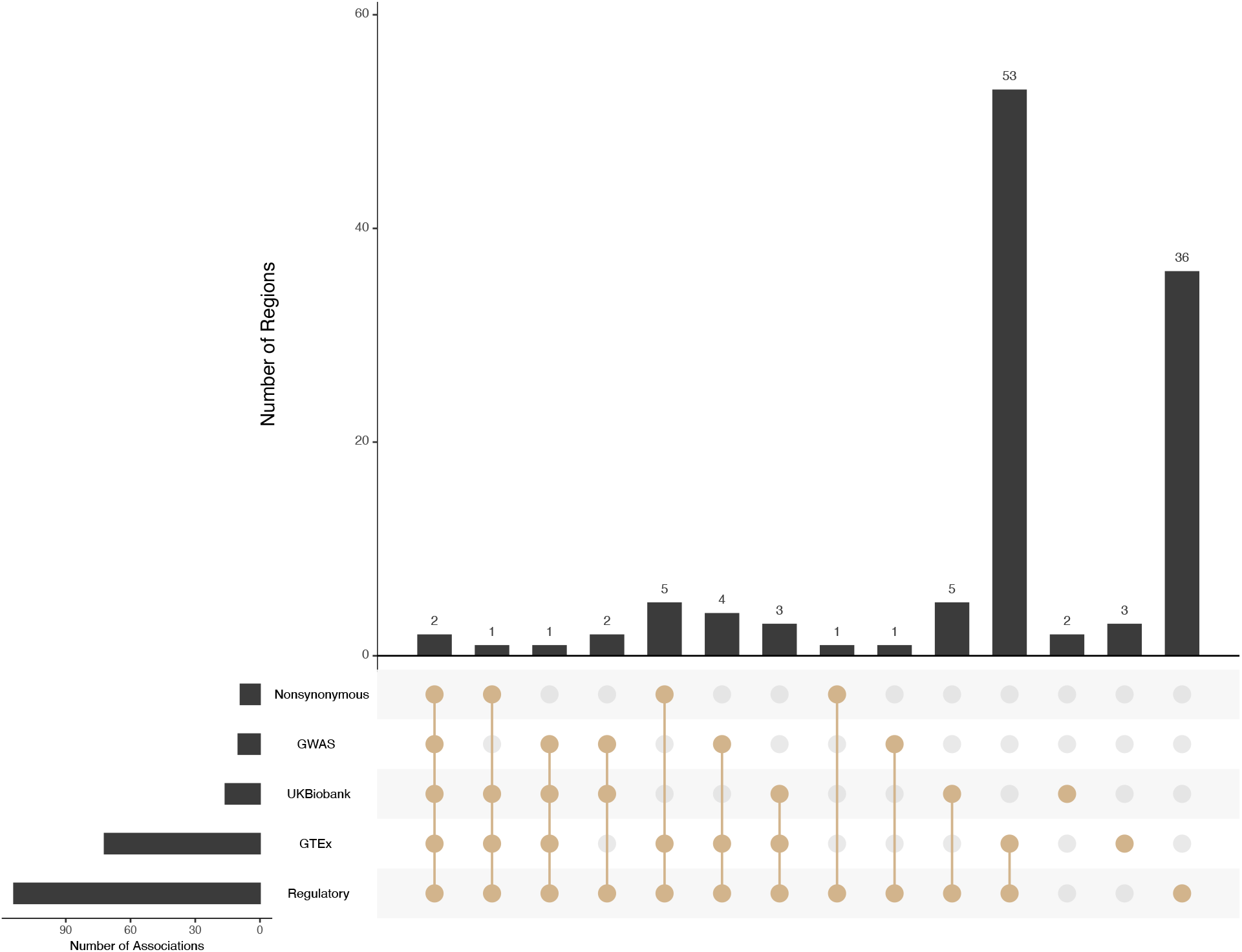
Functional annotations available for the expanded TSP regions. Summary of the annotations of each type available for TSP regions, including tagging variants in high LD with TSPs. A total of 119 out of 125 TSP regions contain at least one line of functional evidence. Multiple lines (two or more) are available for 78 regions.

### Evidence of gene regulatory function for TSPs

We hypothesized that many of the non-coding TSPs in our set perform gene regulatory functions. To evaluate this possibility, we intersected the TSPs and variants in high LD with maps of functional regulatory regions from the Ensembl regulatory build (Zerbino, Wilder, Johnson, Juettemann, & Flicek, 2015). We found 58 TSPs with regulatory annotations in 40 TSP regions, and additionally 1334 LD variants in 114 regions. These include variants in CTCF binding sites, open chromatin regions, promoter flanking regions, enhancers, promoters, and known TF binding sites. Supplementary table 1

Overlap of a variant with a regulatory annotation does not necessarily imply a regulatory function. To consider additional evidence of regulatory function, we examined eQTL in 50 GTEx tissues for overlap with TSPs. At least one eQTL was found for 51 of the TSP regions (40%). Among these 51 regions, 29 TSPs are themselves eQTL; for the remainder, variants in high LD with TSPs were eQTL. The eQTL were found across diverse tissues (Figure 3A), and there was no enrichment for specific gene ontology (GO) terms among the set of genes influenced by TSP eQTL. This suggests that the targets of balancing selection have functions in gene regulation across diverse tissues beyond the immune system.

**Figure 3.**
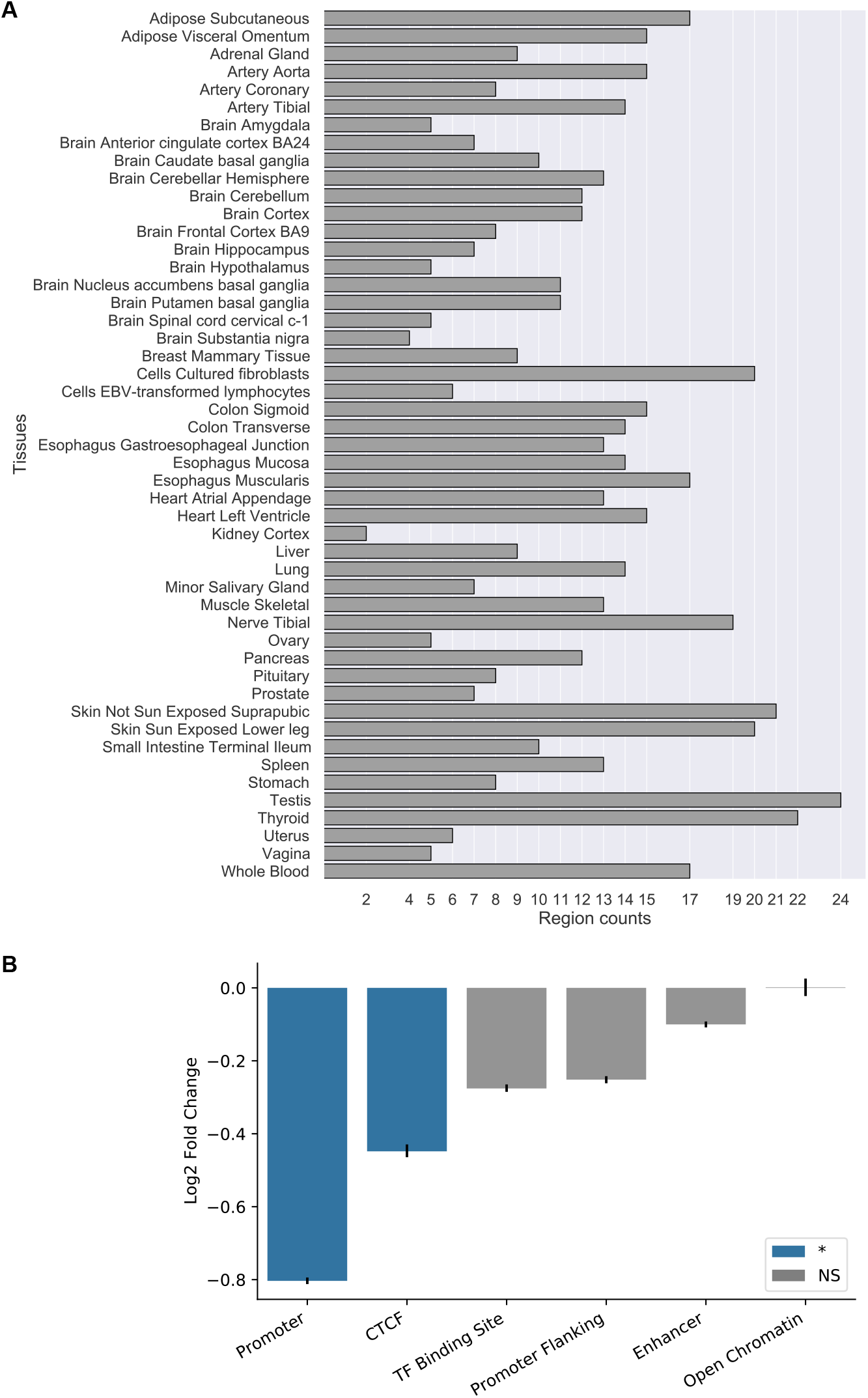
TSPs are eQTLs in diverse tissues and are depleted for overlap with promoters and CTCF sites. (A) The number of TSP regions that contain an eQTL for each GTEx tissue. Variation in TSP regions associates with gene expression in diverse tissues. The associated genes also have Gene Ontology (GO) annotations from diverse functional categories (Supplementary Figure 2). (B) TSP regions are significantly depleted for overlap with promoters and CTCF sites compared to length- and chromosome-matched non-coding regions from the genomic background. The error bars represent 95% confidence intervals.

Next, we tested if the TSP regions are enriched for overlap with any specific types of regulatory regions compared to the expected overlap if they were randomly distributed across the genome. We shuffled the TSP regions 1,000 times maintaining their length and chromosome distributions and avoiding genome assembly gaps and ENCODE blacklist regions and counted the number of overlaps observed for each random permutation. The TSPs overlap slightly fewer base pairs annotated with regulatory functions than expected by chance, with significant (P < 0.05) depletion for promoter and CTCF sites (Figure 3B). Since variants in these regions are likely to influence gene regulation in many tissues (e.g., compared to enhancers which are often context-specific), this suggests that individual TSPs may be less pleiotropic than expected by chance.

### Genome-wide association studies link TSPs to traits

Genome-wide association studies have identified thousands of associations between genetic variants and human traits. We intersected the TSP regions with associations reported in the GWAS Catalog, which is currently composed of 227,262 associations. Since TSPs themselves were not always directly tested in GWAS studies, we also include genome-wide significant (p < 1E-8) associations with the tag variants in high LD with TSPs. We found significant associations for 29 different variants (Figure 4; Supplementary Table 2). Two main functional categories were identified in the GWAS associations for these variants: immunological functions and neurological/behavioral traits. The associations with immune traits were expected given the results of previous balancing selection studies and the few well-characterized instances of LTBS. We identified many variants in LD with TSPs as associated with blood measurement phenotypes and diseases related to immune response failure (chronic inflammatory diseases and ulcerative colitis). We also found many neurological/behavioral related associations among these variants. These traits include cognitive performance (*PLCL1*), chronotype (*RP11-497E19.2*), addiction (alcohol use disorder, *GPR139*; smoking status, *PLCL1*), risky behavior (speeding propensity, *C3orf58*), and mood swings (*PLCL1*). In addition to the immune response and neurological categories, we observed associations with polycystic ovary syndrome, urate levels (*HNF4G*), pancreatic cancer, hepatocyte growth factor levels, and gut microbiota (*OTOS*). We discuss several of these associations in more detail in following sections.

**Figure 4.**
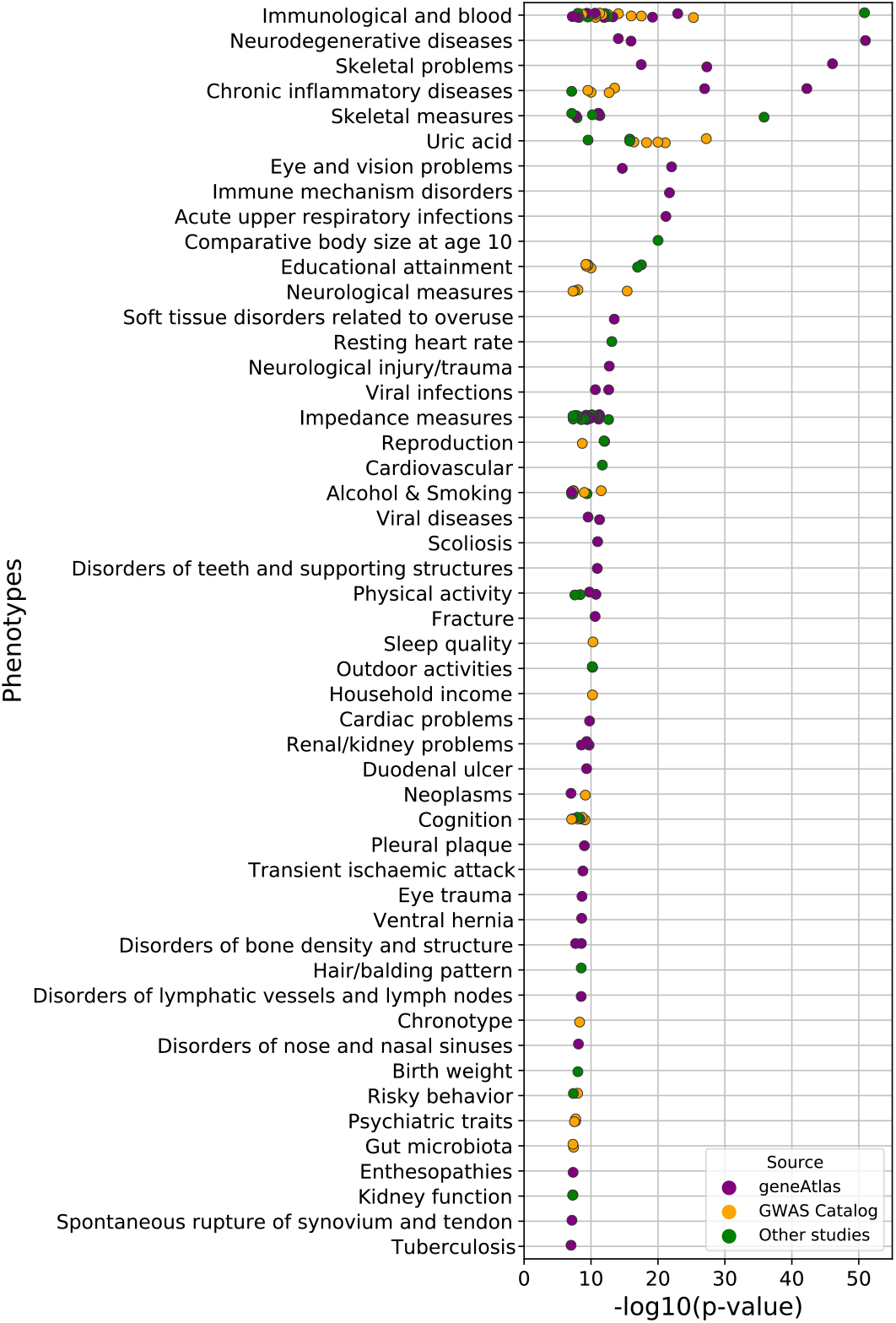
Genome- and phenome-wide association studies link TSPs to diverse traits. Genome-wide significant (P < 1E-8) associations from the GWAS Catalog (yellow) and from a PheWAS over the UK Biobank from the geneAtlas (purple). Each dot represents an association between a TSP region and a trait. Many immune-related traits are associated with TSPs, but there are also associations with a wider variety of phenotypes including osseous, neurological, and nervous system traits. Five extreme associations with immunological and blood traits (P < 1E-60) were truncated for this visualization. Since few TSPs themselves were directly tested in the GWAS Catalog, we include GWAS Catalog associations with tag variants in high LD (r^2^ > 0.8) with TSPs. We also include results from other studies that published genome-wide results using different cohorts.

### Phenome-wide association studies link TSPs to additional diverse traits

The growth of biobanks with linked genetic and phenotypic data has enabled the testing of the association of genetic variants with diverse traits within a single cohort. This PheWAS approach enables exploration of the functional and potentially pleiotropic effects of variants of interest (Bush, Oetjens, & Crawford, 2016). Using published associations from the UK Biobank, we analyzed the association of TSPs with 778 traits; all 125 of the TSP regions were tested. Overall, 21 TSPs in 16 regions had at least one genome-wide significant association (Figure 4, P < 1E-8). Though testing different phenotypes than the GWAS, these associations were qualitatively similar to the GWAS results, in that blood and immune system phenotypes had many associations with TSPs, but the TSPs were also associated with a more diverse set of phenotypes. We found associations in three major categories: immune response traits, body and physical measurements, and neuropsychiatric traits related to addiction. We also observed associations with many other phenotypes, for example: walking habits, heart disorders, renal and kidney problems, pleural plaques, transient ischemic attack, cancerous tumor, and rupture of synovium and tendon (Supplementary Table 3).

### Evidence of protein-coding function for TSPs

The TSPs were originally filtered by Leffler et al. (2013) to be non-coding; however, three of the 263 TSPs are coding variants: a variant containing a synonymous allele in the gene dynein (DNHD1) and another variant that results in a non-synonymous change in a transcript of DNHD1’s paralog DNAH9, and one non-synonymous variant in PKD1L2. This discrepancy is due to changes in genomic annotations over time. For example, PKD1L2 is a calcium channel with potential roles in kidney function. This gene is a polymorphic pseudogene in humans, with the reference genome encoding the pseudogene form; this likely explains why this coding variant was not identified as coding in previous studies. With the LD-expanded set, a total of 9 regions had non-synonymous codon changes (19 variants, Supplementary Table 5). Some of the genes containing these variants include: Sperm Associated Antigen 16 (*SPAG16*), which codes for two proteins that associate with the axoneme of sperm cells; Hepatocyte Nuclear Factor 4 Gamma (*HNF4G*) codes for a receptor involved in DNA binding transcription activity; Leucine Rich Colipase Like 1 (*LRCOL1*) which has enzyme activation activity and is involved in digestion. All of the coding variants had CADD scores suggestive of deleterious effects (scaled C-score > 20). Further work is needed to determine if these coding variants in high LD with the TSPs influence selection.

### Illustrative examples of diverse functions associated with TSP regions

Integrating the above data, we found 79 TSP regions with two or more lines of functional evidence. This included twelve regions with annotations from at least four sources. To illustrate the diverse functions associated with TSPs, we highlight four of these regions.

#### Body mass and alcohol intake

A TSP (rs57790054) on 16p12.3 (hg38.chr16:19992138-20043254) is strongly associated with several growth and body mass phenotypes as well as alcohol intake frequency (Figure 5A; P < 4E-10 for all). Another variant in high LD in Europeans (rs72771074, R^2^=0.89) with a TSP (rs57790054) in this locus was associated with alcohol use disorder in a previous GWAS in a European cohort (P = 5E-8) (Sanchez-Roige et al., 2019). The nearest gene, *GPR139*, encodes for a G-protein coupled receptor expressed in the brain that is involved in alcohol drinking and withdrawal symptoms in rats (Kononoff et al., 2018). This region contains several variants in LD with TSPs in regulatory regions, such as CTCF binding sites (rs117293173, rs13338055, rs74011247, and rs79521770). One TSP (rs57790054) is an eQTL for the gene *KNOP1*, and 26 other LD SNPs are eQTL for both *KNOP1* and *IQCK*. These results suggest that effects on growth and BMI or on addictive behaviors could be under LTBS. We note that there is some evidence of ethanol consumption in chimpanzees, but it is unclear how widespread its availability was over the past several million years (Hockings et al., 2015).

**Figure 5.**
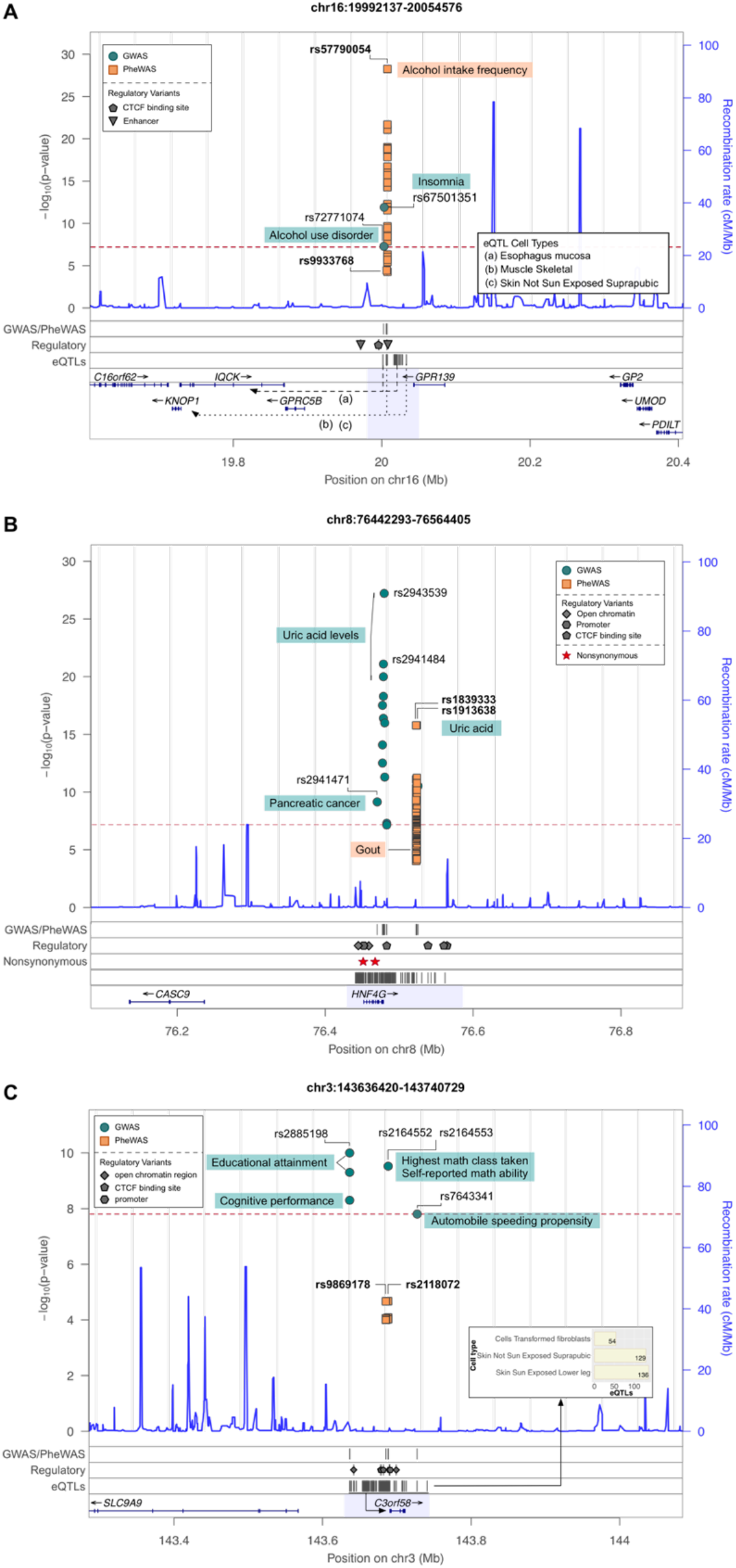
Illustrative examples of non-immune functions associated with TSPs. (A) A TSP in 16p12.3 is associated with body mass index (BMI) and alcohol intake. Regional association plot showing statistically significant genome- and phenome-wide associations (threshold p ≤ 1E-08), regulatory annotations from Ensembl, and eQTLs from GTEx. One of the TSPs (rs57790054, orange) is associated with alcohol intake and several growth and body mass phenotypes in the UK Biobank. A variant in high LD (rs72771074, green) has been associated with alcohol use disorder in a previous GWAS. The TSP is also strongly associated with growth (comparative body size at age 10, 9.6e-21) and body mass index (3.5e-12). The TSPs are nearby *GPR139*, a gene encoding a G-protein coupled receptor expressed in the brain, whose expression levels influence alcohol drinking behavior in rats. The TSP region also contains several CTCF binding sites. TSP are shown in bold text. (B) TSPs in 8q21.11 are associated with urate levels. Regional association plot showing statistically significant genome- and phenome-wide associations (P ≤ 1E-08), eQTL, regulatory and coding SNPs. LD SNPs in this region are associated with urate levels and pancreatic cancer. A TSP (rs1839333) is also associated with gout, although the p-value did not meet our strict threshold. TSP are shown in bold text. TSP are shown in bold text. Figures created using LocusZoom (Pruim et al., 2011). (C) TSP locus on 3q24. Regional association plot showing statistically significant genome- and phenome-wide associations (threshold p ≤ E-08), regulatory and eQTLs. This locus is characterized by neurological traits involved in educational attainment, cognitive performance, and risky behavior (automobile speeding propensity).

#### Urate levels

The TSPs (rs1839333, rs1913638) on 8q21.13 are both significantly associated (P < 2.0e-18) with uric acid levels in multiple GWAS in European and Asian ancestry populations (Figure 5B) (Kanai et al., 2018; Köttgen et al., 2013; Tin et al., 2019). These variants are also associated with a range of body mass traits in the UK Biobank. Another variant in this locus (rs2941471, R^2^=0.97 and R^2^=0.82 in East Asians and Europeans respectively) is associated with pancreatic cancer (p=7E-10). Though elevated uric acid in the blood is associated with many conditions, it is a marker for pancreatic cancer (Stotz et al., 2014). This locus also contains two non-synonymous variants in the gene *HNF4G*, a transcription factor expressed in the liver, kidney, and pancreas, in high LD with the TSPs (rs1805098 and rs2943549). The TSPs are also expression and splicing QTL for HNF4G (P = 1.9E-4 and 8.6E-6, respectively). Variants in *HNF4G* are associated with several traits, including the development of hyperuricemia (Chen et al., 2017).

#### Chronotype

The TSPs (rs1887944, rs7147645) on 14q31.2 are both nominally associated (P <= 9.5e-6) with chronotype in a study of nearly 500,000 people (Jones et al., 2019). This is the only phenotype association with these variants. One variant in high LD (rs17119051) overlaps a regulatory region, but there are no eQTL among the TSPs or other LD variants. Asynchrony in chronotypes in a population has been proposed to be beneficial due to potential protection against predators and other dangers during the vulnerable hours of sleep. This so-called sentinel hypothesis developed out of the observation in Hadza hunter–gatherers of Tanzania that due to variation in sleep phase timing all members of a group are rarely asleep simultaneously (Samson, Crittenden, Mabulla, Mabulla, & Nunn, 2017). Thus, it is possible that LTBS maintained variation at this locus to promote chronotype diversity to mitigate the risks of sleeping.

#### Risky behavior and educational attainment

TSPs (rs9869178 and rs2118072) in 3q24 (hg19.chr3:143636420-143740729) are associated with a range of risky behaviors and educational attainment in both individual GWAS studies and the UK Biobank (Figure 5C). For example, they are associated with automobile speeding propensity (P = 4.4E-8) (Linnér, 2019) and with educational attainment (P = 2.4E-7) (Lee et al., 2018). The TSPs are also modestly associated with variation in brain white matter microstructure (Anterior corona radiata mean diusivities, P = 1.96E-6) (Zhao et al., 2019). Many of the variants in high LD with the TSPs in this region have similar associations and overlap annotated regulatory regions: regulatory open chromatin (rs10662845 and rs6766439), promoter (rs1898263, rs1992094, rs4431106, rs7631704, rs7651567, and rs7653431), and CTCF binding sites (rs7628282, rs7650239, rs7650332, rs9840519, rs9840971, rs9878070, and *rs9840157*). Furthermore, the TSPs (and 159 high LD variants) are significant eQTLs (P ≤ 1E-5) for the gene *DIPK2A* (*C3orf58*) across four GTEx tissues (small intestine terminal ileum, transformed fibroblasts, skin from the lower leg, and suprapubic skin). DIPK2A has not been comprehensively functionally characterized, but it contains a protein kinase domain and is broadly expressed, including in the developing and adult brain. Deletion of this gene has been linked to autism, and its expression is responsive to neuronal activity (Morrow et al., 2008). Associations with behavioral and cognitive traits must be interpreted with caution as these traits are very challenging to quantify and strongly influenced by social factors that may vary with other characteristics. Nonetheless, these associations point to an influence of the TSPs on behaviors relevant to risk tolerance. Thus, it is possible that maintaining a diversity of risk tolerance in human and chimpanzee populations has been beneficial.

## Discussion

In this study we aimed to characterize the function of TSPs with evidence of LTBS. These variants have a deep ancestry in the common ancestor between humans and chimpanzees, and have persisted in the genomes of both species for millions of years. Due to the persistence of these polymorphisms over such long periods, they are likely under balancing selection and have functional effects that influence fitness (De Filippo et al., 2016), yet the majority of the TSPs previously identified do not have known functions. To address this challenge, we identified functional annotations for 114 out of 125 uncharacterized TSP regions with the help of newly developed genomic annotation tools (Figure 2).

Our results show that TSPs likely maintained by LTBS have diverse functions beyond enabling a flexible immune response to pathogens. This is consistent with several recent studies of balancing selection over shorter timescales that have identified regions with functions outside the immune system (Bitarello et al., 2018; Sato & Kawata, 2018; Siewert & Voight, 2017; Viscardi et al., 2018).

The associations we identify suggest possible behavioral, neurological, and other traits that may have driven LTBS. In particular, our results provide support and candidate loci for previous hypotheses about the need for neurological and behavioral diversity in populations. For example, the chronotype association supports the sentinel hypothesis that variability in sleep patterns is the result of natural selection acting to reduce vulnerability of groups while members sleep at night (Samson et al., 2017). Similarly, selection has recently been shown to act on risk-taking behavior in anole lizards (Lapiedra, Schoener, Leal, Losos, & Kolbe, 2018). Thus, our identification of an association between a TSP and human risk-taking behavior (Figure 4C) suggests that LTBS may have maintained genetic variants that contribute to variation in risk taking behavior in humans and chimpanzees. A gene with evidence of eQTL in this region (*C3orf58*) encodes for a protein kinase and has been associated with autism and other neurological disorders (Dudkiewicz, Lenart, & Pawłowski, 2013), adding further evidence of possible neurological drivers to LTBS.

Our results also raise the intriguing possibility that variants that modulate urate levels have been under LTBS. Uricase, the enzyme that metabolizes uric acid into an easily excreted water-soluble form in most mammals, has been lost in great apes. This gene was disabled by a series of mutations that slowly decreased activity over primate evolution, increasing the levels of uric acid in blood (Kratzer et al., 2014). It has been hypothesized that this loss of uricase activity was driven by increase fructose in primate diets due to fruit eating (Johnson et al., 2009; Kratzer et al., 2014). It has also been proposed that high levels of uric acid, a potent antioxidant, played an important role in the evolution of intelligence, acting as antioxidant in the brain (Álvarez-Lario & Macarrón-Vicente, 2010). However, as reflected in the associations with this locus, elevated uric acid levels contribute to many common diseases in modern humans, including chronic hypertension, cardiovascular disease, kidney and liver diseases, metabolic syndrome, diabetes, and obesity (Gustafsson & Unwin, 2013). This suggests potential functional tradeoffs at this locus. However, we emphasize that proving the environmental drivers of past selection is challenging.

Some of the phenotype associations we discovered may reflect manifestations of variation on traits in modern environments that could not be long-term drivers of balancing selection. As an extreme example, influence on smoking behavior could not have been the cause of LTBS given the relatively recent wide availability of alcohol and nicotine. Though we note that there is some evidence of ethanol consumption in chimpanzees (Hockings et al., 2015). Even if they reflect modern environments, these associations provide hints about possible behavioral, neurological, or other traits that may have driven LTBS. For instance, plant chemicals can hijack reward systems in the brain that motivate repetition and learning (U.S. Department of Health & Human Services, 2016). The same systems that influence these action and consequently reproductive fitness potentially created a byproduct of excessive seeking of dopamine or other reward chemicals.

There are several caveats to our work. First, even with recent growth of genetic and phenotypic databases, our knowledge of the functions of most regions of the genome is sparse. Thus, failure to observe a functional association does not imply that a region does not have an important function. Second, the genome- and phenome-wide association tools we used are limited to the samples that have been analyzed; available data do not represent the full scope of human variation. Most of the individuals analyzed in available genetic association studies are of European ancestry. Variant functions and the ability to detect associations vary across human populations; however, we anticipate that TSPs should have functional effects across populations, unless modern environments have masked the pressure driving LTBS. Third, even in PheWAS, a limited number of phenotypes have been quantified across individuals, and these studies are focused on a subset of clinically relevant traits rather evolutionarily relevant traits. Fourth, in some analyses, we considered annotations based on trait associations with variants in high LD (r^2^ > 0.8) with TSPs. This could potentially introduce false positives if the variant also tags a different causal non-TSP variant. Given the long-term selection on TSPs, they are strong candidates for casual variants, but functional studies are needed to confirm these statistical associations. Finally, our analyses have focused on the human context; due to lack of functional data, it is not possible to explore the function of TSPs in chimpanzees. Nonetheless, we feel that our integration of genome-scale annotations and biobank data highlight the diversity of functions associated with LTBS.

In conclusion, we assign putative functions for TSP regions that likely persisted due to balancing selection dating back to at least the common ancestor of humans and chimpanzees. These annotations expand beyond immune functions to traits relevant to behavior, cognition, and body shape. Notably, we also find that most regions with multiple TSPs overlap gene regulatory annotations suggesting LTBS on gene expression levels. As methods improve for quantifying the effects of variants on gene regulation in different tissues and how these relate to organism-level phenotypes, we anticipate deeper mechanistic understanding of the functions and potential evolutionary pressures on these regions.

## Methods

### Trans-species polymorphisms

The initial set of 263 TSPs from 125 regions analyzed in this study was published by Leffler et al. (2013). The set is composed of trans-species polymorphisms that are shared between 51 Yoruba individuals in the 1000 Genomes Pilot 1 and 10 chimpanzees from the PanMap project. Each SNP is part of a region composed of two or more TSPs within 4 kb of each other, where the TMRCA between polymorphisms is deeper than the speciation event between humans and chimpanzees.

To increase our ability to identify annotations in each locus, the dataset was expanded to include variants in high LD (threshold R^2^=0.8) with each of the TSPs as is common in association studies. We computed linkage disequilibrium with the TSP variants from 1000 Genomes Project Phase 3 data using rAggr a web tool developed by the University of Southern California (http://raggr.usc.edu). We considered LD in African, East Asia, and European populations. Variants with no reported RSID were excluded from the analysis. The dataset was thus expanded by 9,996 SNPs in high LD with the TSPs for a total of 10,259 SNPs.

### Genome- and Phenome-wide associations

The GWAS Catalog collects variant-trait associations from published genome-wide association studies. The database is currently composed of more than 200,000 associations. We used the GWAS Catalog version October 2020 to find functional associations for the LTBS variants. The search was done using the BEDTools intersect function between the GWAS catalog and the LD-expanded TSP dataset (Quinlan, 2014).

PheWAS is an analysis strategy built on top of medical records with information about patient phenotypes and associated variants. The geneAtlas catalog (http://geneatlas.roslin.ed.ac.uk/phewas/) takes advantage of the data provided by the UK Biobank cohort, which contains medically relevant data from nearly 500,000 British individuals of European ancestry. This database contains 3 million variants in 778 traits. We matched our set of variants against the geneAtlas database to search for traits associated with LTBS.

### GTEx eQTL data

To evaluate potential gene regulatory effects of TSPs in non-coding regions, we analyzed data from GTEx, a project developed to quantify the consequence of genetic variation on expression at the tissue level (https://www.gtexportal.org/home/). The GTEx project v8 data have identified eQTL across 50 tissues based on analyses of nearly 1,000 individuals to identify differential expression through SNP variation. The intersection between the TSPs and LD SNPs and the GTEx eQTL returned a large collection of TSP eQTL.

### Other functional categories

We used ANNOVAR to identify variants overlapping protein coding regions. We used the Ensembl Regulatory Build (Zerbino et al., 2015) to identify variants overlapping regions with regulatory function. In all, 58 TSPs in 40 regions and 1334 LD SNPs in 114 regions overlapped regulatory annotations.

### Enrichment for overlap with regulatory regions

We used a permutation testing framework to calculate whether TSPs were more enriched for overlap with regulatory regions than expected by chance (Benton, Talipineni, Kostka, & Capra, 2019). We quantified the number of overlapping TSPs for each type of regulatory region (open chromatin, promoter, enhancer, promoter-flanking, CTCF binding site, TF binding site). We then compared the observed TSP overlap to a null distribution of expected overlap generated by randomly shuffling the regulatory regions 1000 times and obtained an empirical p-value. We maintain the original length distribution and chromosome for shuffled regions and exclude all ENCODE blacklist and gap regions (Kundaje, 2013).

## Declarations

### Data Availability Statement

The data underlying this article are available in the article and in its online supplementary material.

### Author’s contributions

Conceptualization: JAC; Methodology: KV, MLB, JAC; Investigation: KV, MLB, JAC; Writing – Original Draft: KV, JAC; Writing – Review & Editing: KV, MLB, JAC; Funding Acquisition: JAC; Resources: JAC; Supervision: JAC.

## Acknowledgements

We thank Evonne McArthur, David Rinker, and other members of the Capra Lab for helpful comments on this work. This work was conducted in part using the resources of the Advanced Computing Center for Research and Education at Vanderbilt University, Nashville, TN.

