## Supplemental Figures for "Diverse functions associate with trans-species polymorphisms in humans"

### Supplementary Figures

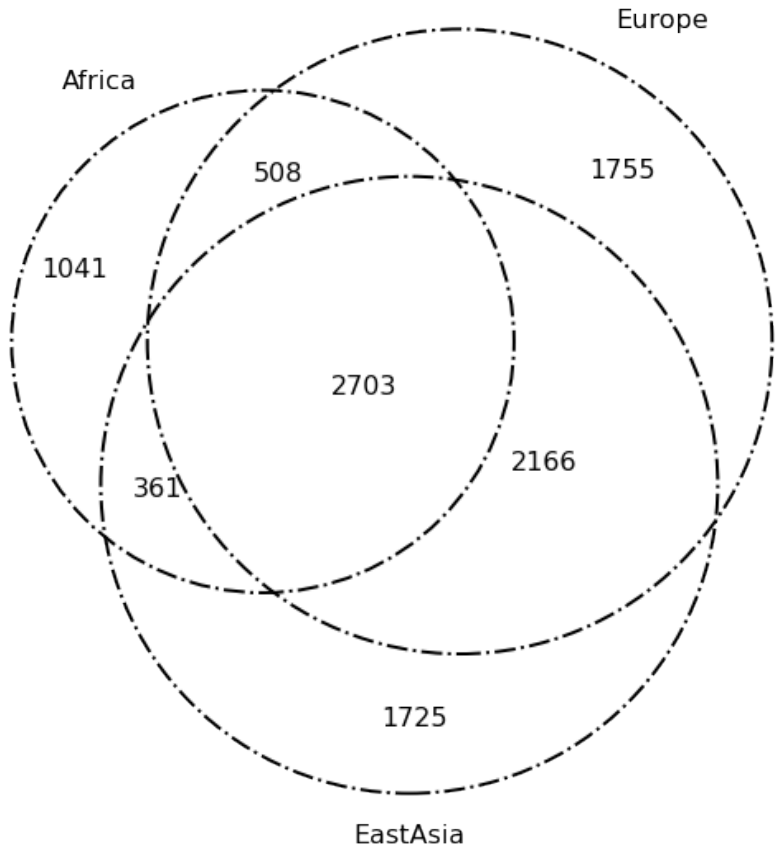


**Supplementary Figure 1. Trans-species polymorphisms (TSPs) likely resulting from long-term balancing selection (LTBS).** We consider 125 regions containing 263 TSPs. Two cases are possible for these haplotypes: one in which one site is under selective pressure while the other is neutral, and one in which the shared sites have epistatic functions and are both under selection. In addition to the 263 TSPs, we also considered functional associations with 10,259 variants in high LD ($r^2 > 0.8$) with a TSP at least one population from the 1000 Genomes Project (Supplementary Figure 1).

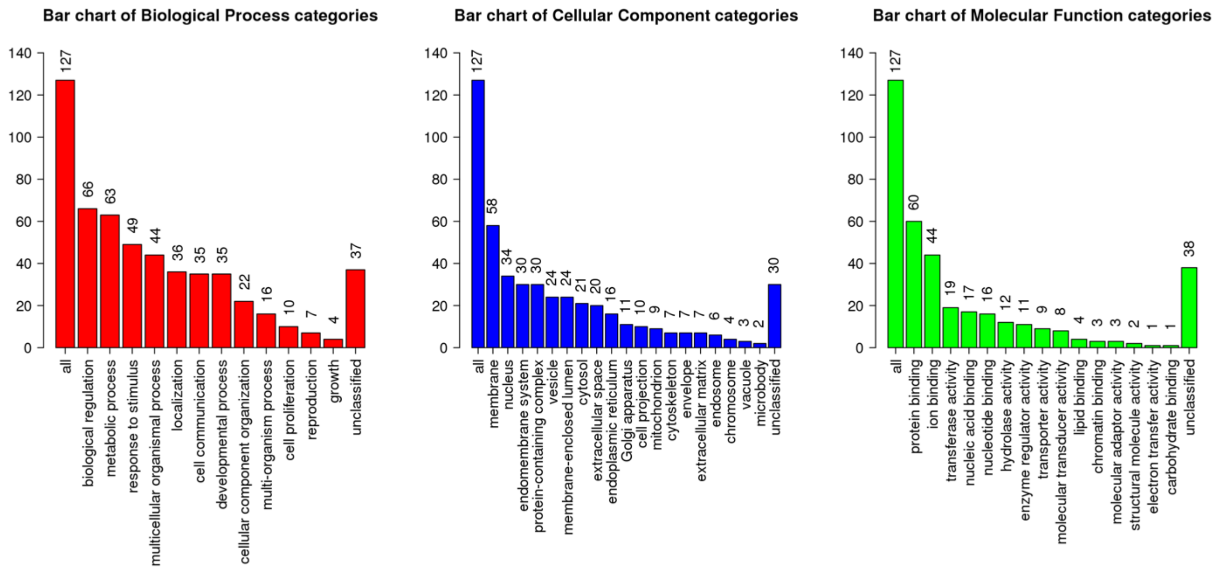


**Supplementary Figure 2. GO analysis of genes with evidence of eQTL.** A Gene Ontology (GO) enrichment analysis shows no significant enrichment on any TSP associated gene set. The analysis was performed using WebGestalt.
